# Reconciling global priorities for conserving biodiversity habitat

**DOI:** 10.1101/850537

**Authors:** K. Mokany, S. Ferrier, T.D. Harwood, C. Ware, M. Di Marco, H.S. Grantham, O. Venter, A.J. Hoskins, J.E.M. Watson

## Abstract

Degradation and loss of natural habitat is the major driver of the current global biodiversity crisis. Most habitat conservation efforts to date have targeted small areas of highly threatened habitat, but emerging debate suggests retaining large intact natural systems may be just as important. We reconcile these perspectives by integrating fine-resolution global data on habitat condition and species assemblage turnover, to identify Earth’s high-value biodiversity habitat. These are areas in better condition than most other locations once supporting a similar assemblage of species, and are found within both intact regions and human dominated landscapes. However, only 18.6 % of this high-value habitat is currently protected globally. Averting permanent biodiversity loss requires clear spatially explicit targets for retaining these unprotected high-value habitats.

---

The world is facing a biodiversity crisis, with up to half a million species committed to extinction over the coming decades (*1–4*). The most important threat to biodiversity is ongoing degradation and loss of natural habitat (*4–6*), with more than half of the world’s land surface now under human-dominated land uses (*7*). Retaining remaining natural habitat is a crucial response to limiting further extinctions, with recent proposals for post-2020 protected area targets of 30 % of the planet by 2030 (*8*), and more ambitious calls for protection of half the terrestrial biosphere by 2050 (*9–11*).

While there is agreement on the need to retain natural habitat to prevent species extinctions, division remains over which areas are most important for focusing the limited resources available for conservation. Brooks and colleagues (2006) first characterized two contrasting general strategies to prioritizing habitat retention: ‘proactive’ approaches prioritize retention of the most intact natural systems, such as large areas of wilderness (*12–14*), whereas ‘reactive’ approaches highlight the benefits of protecting and managing smaller areas within more degraded regions, where many threatened species occur (*15–17*). Reconciling these differing perspectives is crucial in identifying which areas we need to most urgently protect and manage in order to best promote the persistence of biodiversity globally (*18*). This is particularly important in the light of discussions around a post-2020 biodiversity framework under the Convention on Biological Diversity (CBD) (*19*).

Proactive and reactive habitat conservation strategies diverge in how they consider habitat condition – the degree to which a location has been impacted by human activities, such as vegetation disturbance, hunting, or invasive alien species. Proactive strategies focus primarily on the habitat condition of each location, prioritizing large areas that are in the best condition (*12–14*). In contrast, reactive strategies focus on the condition of habitat across the entire ranges of species, communities or ecosystem types, providing an indication of the conservation status of those biodiversity elements and the vulnerability of individual locations to future degradation (*15–17*).

Here we present an analytical framework that reconciles these differences to help identify priority areas around the world where protection and management will best promote biodiversity persistence. Our approach effectively integrates both the condition of a location itself and the condition of all other locations expected to have supported shared species prior to any habitat degradation. To reconcile proactive and reactive strategies for identifying high-value biodiversity habitat, we analyze where the condition of a given location sits relative to the frequency distribution of condition levels for all biologically-similar locations – a property we call “contextual intactness”.

Our analysis combines fine-resolution (1 km) global datasets on habitat condition and spatial biodiversity patterns (*20*). We apply an updated map of the terrestrial human footprint on natural systems to describe current habitat condition (Fig. S1) (*21*). To inform the biodiversity value of each location, we harness generalized dissimilarity models of species assemblage turnover for terrestrial vertebrates, invertebrates, and plants, derived using more than 100 million occurrence records from more than 400,000 species (*22, 23*). These models enable the estimation of the similarity in species assemblages between any pair of locations, and subsequently the expected uniqueness of the biodiversity within any terrestrial location globally.

## Results and Discussion

By coupling these two data sources, we assessed the contextual intactness of each location (1 km grid cell), by determining the proportion of habitat expected to have once supported a similar assemblage of species but is now in worse condition than the focal location (Fig. 1) (*20*). Locations with the highest contextual intactness are where all other biologically-similar locations are in a worse condition. We found areas of high contextual intactness in every biogeographical realm (Fig. 2), spanning regions with large coverage of wilderness as well as regions with largely degraded landscapes.

**Fig. 1.**
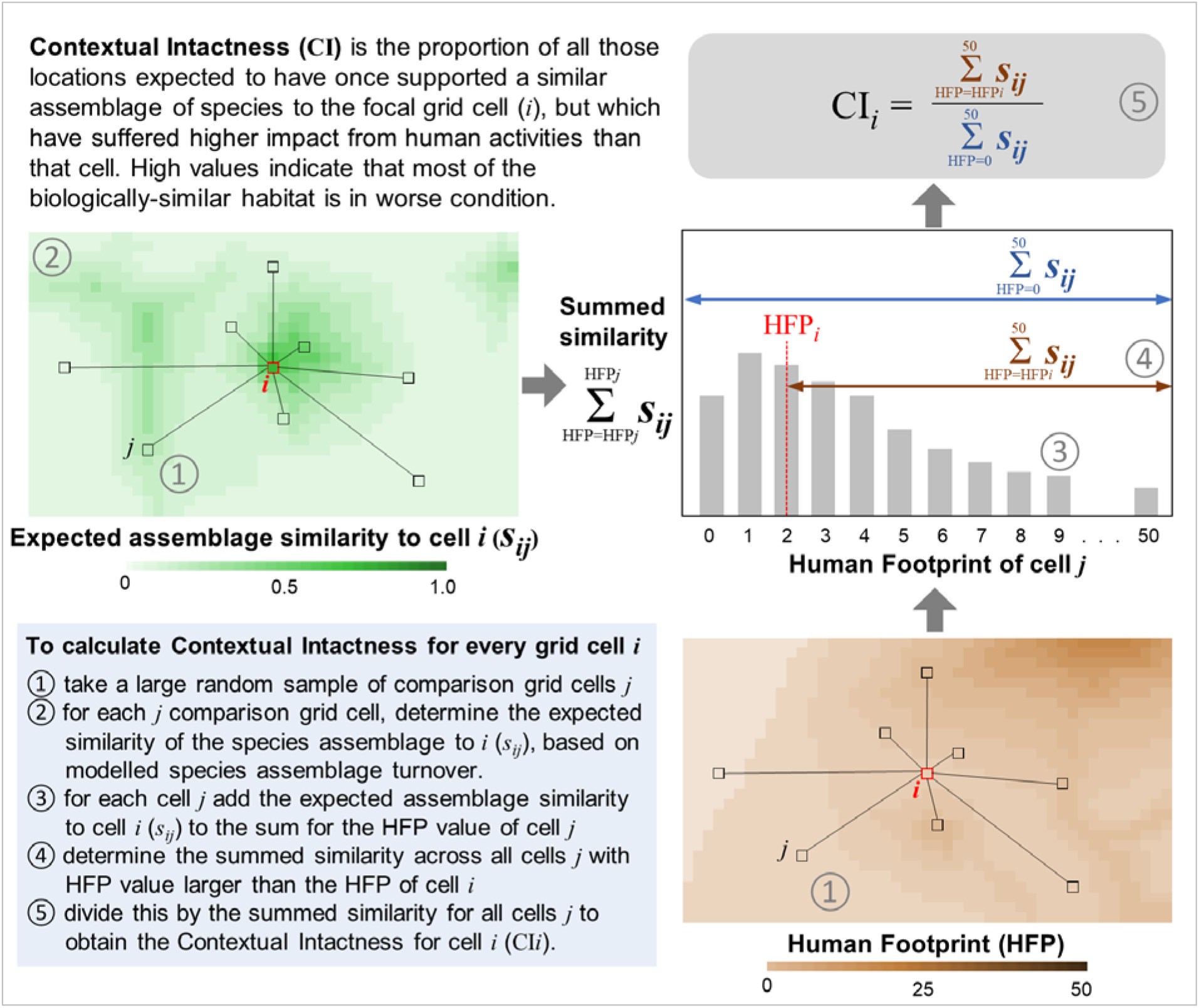
Conceptual depiction of the analytical approach to quantifying the contextual intactness for each terrestrial 1 km grid cell globally.

**Fig. 2.**
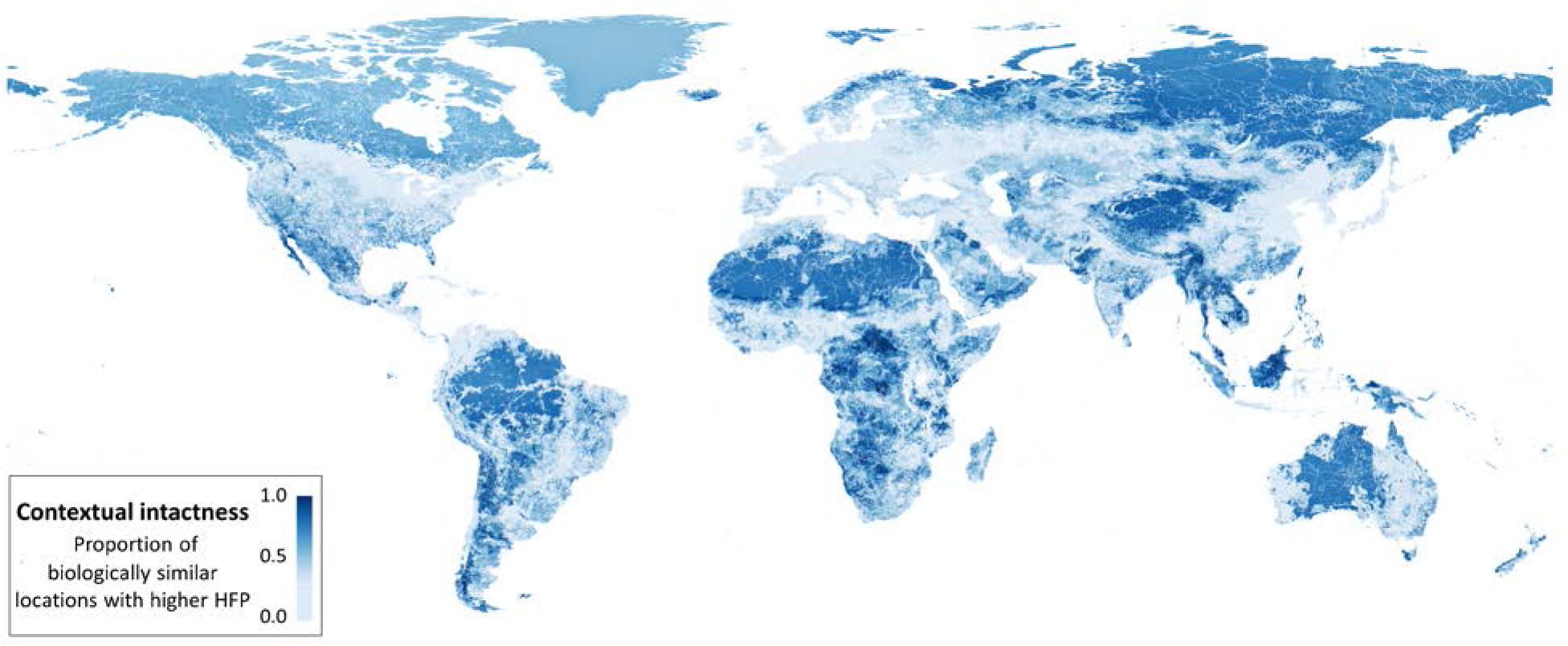
Contextual intactness of habitat for biodiversity. For each location, the proportion of habitat expected to have once supported a similar assemblage of species but is now in worse condition than the focal location (higher human footprint - HFP). The result is averaged across three broad terrestrial taxonomic groups: vertebrates, invertebrates and vascular plants.

Because contextual intactness integrates both habitat condition and patterns in species assemblage turnover, it varies markedly within a given level of human footprint (Fig. 3). For example, habitat at the edge of wilderness areas often has a higher contextual intactness than core areas, as these edge locations are more likely to share species with locations outside of wilderness and subject to higher levels of human impact (Fig. 2). Indeed, our analysis shows that a location with a higher human footprint can still have a higher contextual intactness value, if it is the ‘best-on-offer’ in terms of the condition of all other locations with similar expected species assemblages (Fig. 3). Such areas include remnant habitats within heavily modified regions, such as temperate biomes with extensive agricultural land use. Similarly, while each of the world’s terrestrial biomes contain some habitat with high contextual intactness (fig. S2), the distribution of contextual intactness across biomes does not directly align with the distribution of the human footprint, due to neighboring biomes sharing species in common. For example, temperate coniferous forests have relatively low human footprint, but they share some species with the relatively unimpacted boreal forests (Fig. S2) (*20, 24*).

**Fig. 3.**
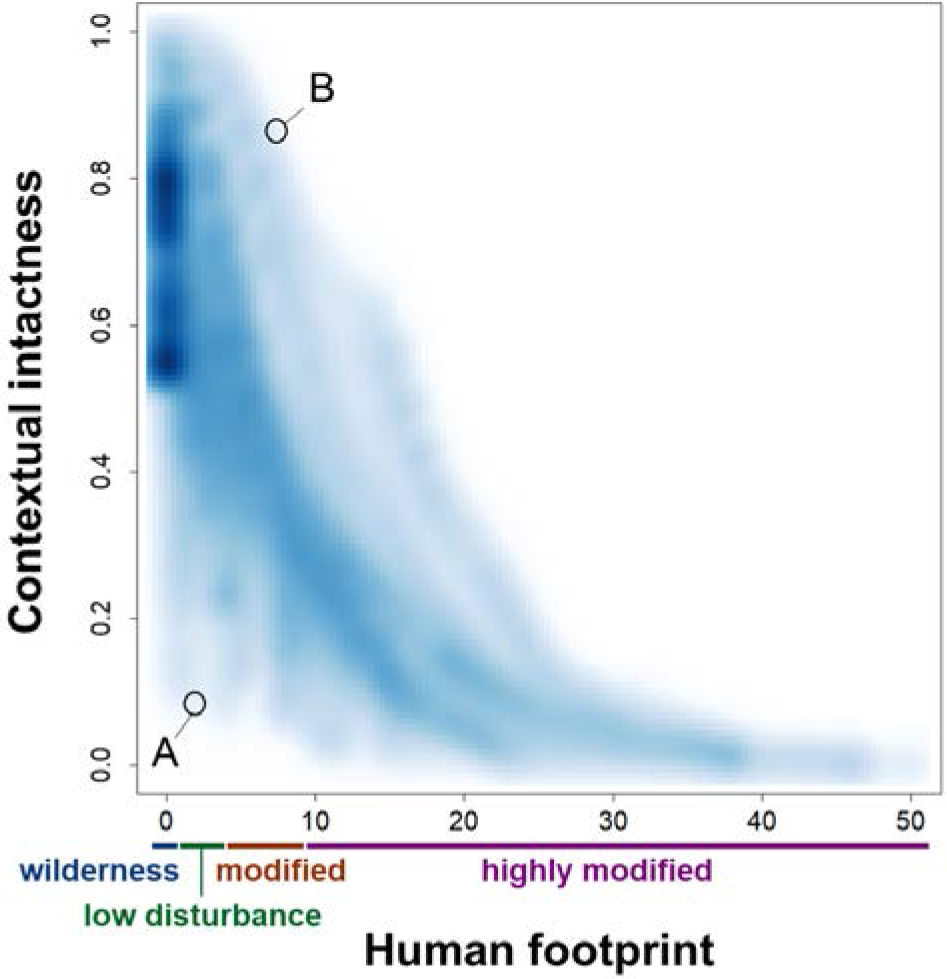
Within a given level of human footprint, there is wide variation in contextual intactness values for terrestrial biodiversity. For example, location A has low levels of disturbance but has a high proportion of similar habitat in wilderness areas. In contrast, location B has a higher level of human footprint, but most of the areas with similar expected species assemblages are in even more highly modified areas. Data are a sample of 430,000 locations (1 km grid cells) shown as color density with kernel bandwidth of 0.75.

Our analysis integrates both proactive and reactive prioritization strategies into one cohesive conservation framework, which we demonstrate by mapping contextual intactness against four categories of the human footprint – ‘wilderness’, ‘low disturbance’, ‘modified’ and ‘highly modified’ (Fig. 4). This shows that while large intact areas are important, partially degraded fragmented habitat can also play a vital role in conserving biodiversity, given that these areas may support species that have lost a large proportion of their habitat elsewhere through human land use (Fig. 4). For example, the tropical forests of south-east Asia have experienced widespread loss (*25, 26*), so even if remaining areas are slightly modified, they are of high-value in sustaining the unique diversity of the region (Fig. 4B) (*27*).

**Fig. 4.**
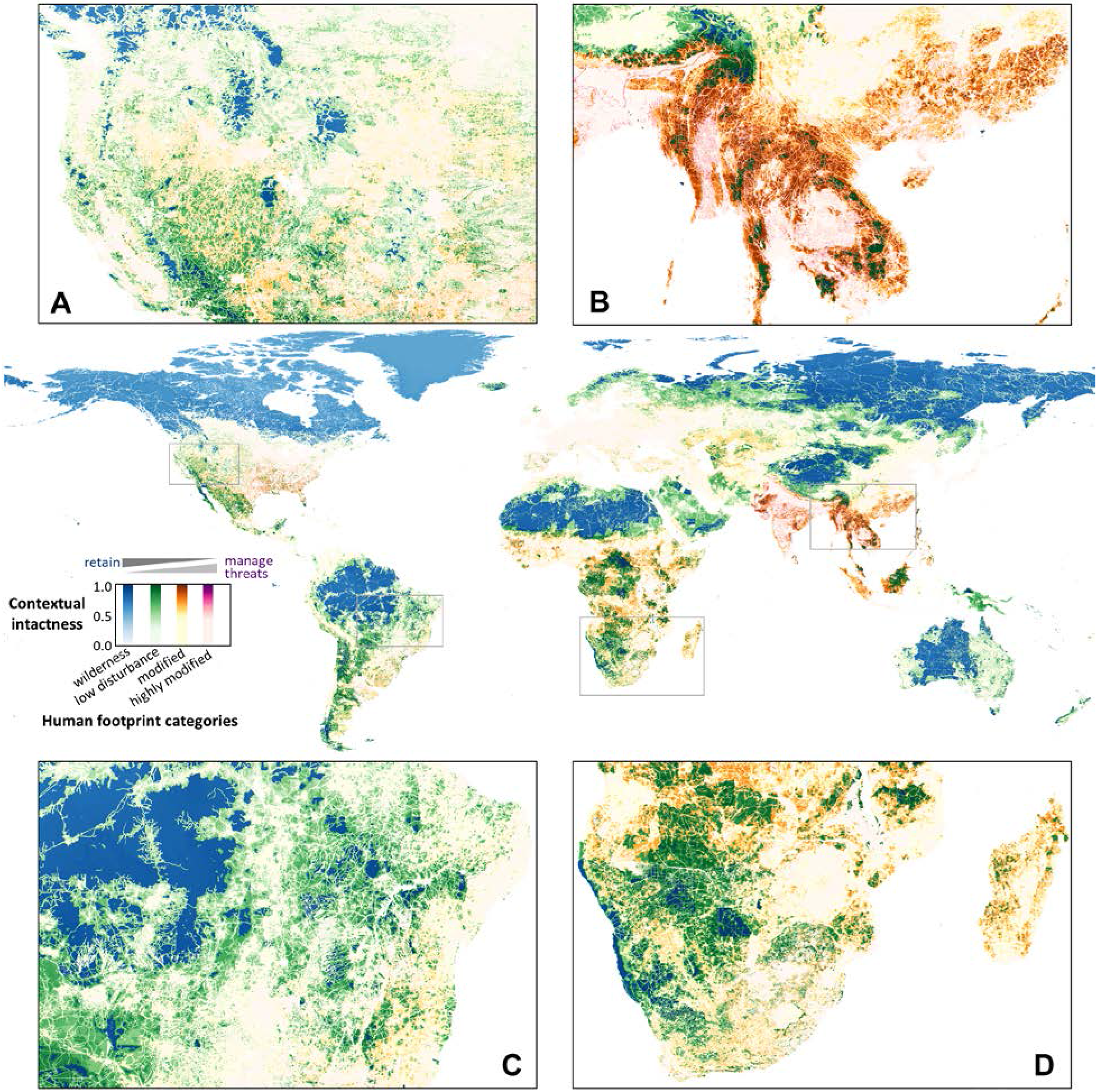
High-value biodiversity habitat across levels of human footprint. Contextual intactness represented across four human footprint (HFP) categories: wilderness (blue, HFP = 0); low disturbance (green, HFP = 1 - 3); modified (orange, HFP = 4 - 9), and highly modified (pink, HFP ≥ 10). Darker shades represent higher contextual intactness. Panels show greater detail for (**A**) north-west North America; (**B**) south-east Asia; (**C**) south-east Amazonia; (**D**) southern Africa.

In addition to the very broad taxonomic scope of our global assessment, a key feature is the spatial resolution at which our analyses are performed (Fig. 4A-D), which is fine compared to many previous global conservation assessments typically implemented at the ecoregion or biome level (*10, 15, 28*). A fine spatial resolution helps account for local impacts of human activities on nature, and the close affinities many species have with environmental and habitat features that can change over very short distances (*29*). While not obvious at the global scale, most heavily impacted regions contain very small areas of high-value habitat that is important for biodiversity conservation (Fig. 4A-D). Our finely resolved assessment thus provides not only a measure of contextual intactness across the land surface of the planet, but does so at a spatial resolution of relevance to conservation policy, planning, and management at the regional or national level. Priority actions for high-value habitat will likely vary from retaining and maintaining current low-disturbance of wilderness areas, to actively managing ongoing threatening processes in more highly modified areas (Fig. 4).

To help inform progress toward renewed global protected area targets (*8, 10, 11*), we identified high-value biodiversity habitat as locations with contextual intactness greater than 0.5. These are locations which are in better condition than more than half of the areas expected to have supported a similar assemblage of species, and hence they cover approximately half the land surface of the planet. We then assessed the degree to which these high-value habitats occur within protected areas, finding that only 18.6 % of the world’s high-value habitat for biodiversity is currently protected (fig. S3) (*20*). Therefore, protected areas are only slightly better than random at safeguarding the highest value habitat for biodiversity around the world, given 14.9 % of Earth’s land surface is protected (*30*). The largest areas of unprotected high-value biodiversity habitat are in wilderness and areas experiencing minimal human disturbance (Fig. S3) (*14*). However, high-value habitat in regions that have been more heavily modified has an even lower proportional coverage in protected areas, reducing to 6 % protected (fig. S3).

There have been decades of debate on whether it is more urgent to preserve intact landscapes or to protect areas facing high human pressure (*12–17*). Our findings demonstrate that this conservation dichotomy is unnecessarily artificial, and that the situation is actually more nuanced, with high-value biodiversity habitat spanning all levels of human pressure. These results show that the contribution of each location to the persistence of biodiversity is entirely dependent on the regional and environmental context, through the spatial patterns in both biodiversity and habitat condition. Preserving these high-value areas should be integral to the international commitments to halt biodiversity loss (*31*), as habitat degradation there would make it much harder to achieve international biodiversity targets.

Retaining high-value habitat for biodiversity also requires supportive measures beyond establishing protected areas, including strong legislation and programs to limit threats to intact habitat (*4*). This includes establishing and policing laws that limit land clearing and forest degradation, in addition to programs that support sustainable socio-economic development outside of high-value habitat areas. Jurisdictions where habitat protection laws and policies have been weakening, rather than strengthening (*32, 33*), need to be encouraged and supported by the global community to identify ways to improve human wellbeing while retaining the habitat required to support the Earth’s biodiversity (*4*).

Given the irreversibility of species extinctions, society must act now to retain the Earth’s unique evolutionary heritage. The high-value habitat we have identified will be crucial for the persistence of biodiversity into the future, requiring strong commitments by governments, businesses and society to stop their loss and degradation.

## Methods

### Broad approach

The broad approach of our analysis was to assess the contextual intactness of every terrestrial location across the globe, by combining data on two features: (i) the current habitat condition of every location, and; (ii) the expected similarity of the species assemblage occurring in each location relative to every other location. To quantify the current habitat condition, we applied a map of the human footprint (*21*), updated to the year 2013 using some revised spatial datasets as described below. To quantify the expected similarity in species assemblages between any pair of locations, we utilized generalized dissimilarity models of species assemblage turnover for terrestrial vertebrates, invertebrates, and plants across the terrestrial surface of the globe (*22*). These models are also described in more detail below. Using these two data sources, we developed and applied a new analysis to derive the contextual intactness across the globe, which for each location is the proportion of habitat that would be expected to host a similar assemblage of species but is in a worse condition (Fig. 1).

### Human footprint

The human footprint is a standardized measure of the cumulative human footprint on the terrestrial environment at 1 km resolution (*21*). The original global human footprint layer was developed by Sanderson *et al*. (*34*) based on data from the early 1990s, considering four types of data as proxies for human influence: (i) human population density; (ii) land transformation; (iii) human access, and; (iv) electrical power infrastructure. Datasets representing these features were weighted and summed to derive a human footprint score ranging from 0 to 72 (*34*).

To provide an updated assessment of human pressures on the environment, Venter *et al*. (*21*) utilized data centered around two timepoints: 1993 and 2009. This analysis harnessed data on eight human pressures: (i) extent of built environments; (ii) crop land; (iii) pasture land; (iv) human population density; (v) night-time lights; (vi) railways; (vii) roads, and; (viii) navigable waterways. These pressures were weighted following the approach of Sanderson *et al*. (*34*), and then summed to create the standardized human footprint for all non-Antarctic land areas, ranging from 0 to 50 (*21, 35*).

Here we apply a revised version of the human footprint assessment undertaken by Venter *et al*. (*21*). The same approach and weighting scheme was used, but harnessing updated datasets centered on the year 2013. This results in the same 0 – 50 range of human footprint scores, but with each location having a value that reflects the more up-to-date impacts of human activities on the natural environment (Fig. S1).

In communicating the results of our analysis, we applied four categories of the human footprint (HFP), based on extensive experience of the co-authors with the human footprint index:

- ‘wilderness’ (HFP = 0);
- ‘low disturbance’ (HFP = 1 - 3);
- ‘modified’ (HFP = 4 - 9);
- ‘highly modified’ (HFP ≥ 10).

### Similarity of species assemblages

To obtain estimates of the similarity in species assemblages between any pair of terrestrial locations, we utilized generalized dissimilarity models derived by Hoskins *et al*. (*22*) in the BILBI modelling framework (Biogeographic Infrastructure for Large-scaled Biodiversity Indicators). Rather than modelling each species individually, this approach models the similarity in the species assemblages between pairs of locations (30-arcsecond grid cells – approximately 1 km) based on the environmental conditions at each location and the geographic distance between them. These models then enable prediction of the expected similarity in species assemblage based on spatially complete environmental surfaces. This approach overcomes biases and deficiencies of raw biological data across different regions of the planet, enabling comprehensive biodiversity assessments from regional to global scales (*22*). While we present an overview of the modelling of pairwise assemblage similarity here, a full description is provided by Hoskins *et al*. (*22*).

The BILBI framework includes models of similarity in species assemblage for three broad taxonomic groups: vertebrates, invertebrates, and vascular plants. For each of these three broad taxonomic groups, species occurrence records were obtained from GBIF then filtered for accuracy. The subsequent modelling utilized 41,082,043 records for vertebrates (across 24,442 species), 13,244,784 records for invertebrates (across 132,761 species), and 52,489,096 records for vascular plants (across 254,145 species).

As predictors of site-pair species assemblage similarity, the BILBI framework utilizes a standard set of 15 environmental variables, including: five soil variables (bare ground, bulk density, clay, pH, silt) (*36*), two terrain variables (topographic roughness index, topographic wetness index) (*37, 38*) and eight topographically adjusted climate variables (annual precipitation, annual minimum temperature, annual maximum temperature, maximum monthly diurnal temperature range, annual actual evaporation, potential evaporation of driest month, maximum and minimum monthly water deficit) (*39, 40*).

Models of species assemblage similarity were fit using a novel extension of generalized dissimilarity modelling (GDM) (*41*), where the response variable is the probability that a pair of species records drawn randomly from two locations represent two different species rather than the same species. This modelled probability is then back-transformed to the common 0 to 1 measure of similarity in ecological communities (similar to the Sørensen index) for prediction and analysis (*22*). For each of the three broad biological groups, individual models were developed for each of the 61 unique bio-realms defined in the WWF’s nested biogeographic-realm, biome and ecoregion framework (*42*), with species records in neighboring bio-realms also used for model fitting. Model performance varied across taxonomic groups and bio-realms, with an average Mann Whitney U statistic of 0.73 for vertebrates, 0.70 for invertebrates and 0.79 for plants.

The models of site-pair similarity in species assemblages were projected spatially as GDM transformed grids, using the environmental predictor layers. The GDM transformed grids enable subsequent prediction of the similarity in species assemblage between any pair of grid cells, hence provide the capacity for a broad range of analyses, rather than providing just a single spatial layer. For example, the BILBI framework has been applied to assess the impacts of global climate and land-use scenarios on plant biodiversity (*23*), the importance of wilderness areas for the global persistence of biodiversity (*14*), and forms the basis for two indicators endorsed by the Convention on Biological Diversity for reporting against Aichi Targets: (i) the Protected Area Representativeness and Connectedness index (PARC), and; (ii) the Biodiversity Habitat Index (BHI) (*43*).

### Quantifying contextual intactness

To determine the contextual intactness for each location across the globe, we combined the revised human footprint spatial layer and the predicted similarity in species assemblages between pairs of locations using the BILBI framework (Fig. 1). For each grid cell *i*, we selected a spatially regular randomly positioned selection of *n* other grid cells *j* to compare to cell *i*. A sample of comparison cells is required because there are >200 million cells on the 1 km terrestrial grid of the planet, and comparing each grid cell with every other grid cell is computationally prohibitive. For this assessment, the number of other grid cells *j* was a minimum of 1% of the total grid cells within each of the world’s 7 biogeographic-realms (Antarctica being excluded) (*42*).

We then determined the expected similarity (*s_ij_*) in species assemblages between cell *i* and each comparison cell *j* using the BILBI framework (*22*). The human footprint (HFP) value for cell *i* (*HFPi*) and all comparison cells *j* (*HFPj*) was also extracted. We then derived a histogram of the summed species assemblage similarity to grid cell *i*, within integer bands of the human footprint value for all the comparison cells 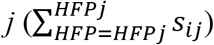. From this histogram, we then calculated: (a) the sum of the assemblage similarities to *i* where the comparison cell *j* had a higher human footprint to *i* 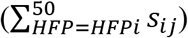, and; (b) the total sum of the all the assemblage similarities between *i* and *j* across all human footprint scores 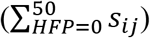. The contextual intactness for grid cell *i* (*CI_i_*) was then calculated as the sum of assemblage similarities to *i* with a higher human footprint divided by the total sum of assemblage similarities to *i* (Fig. 1):

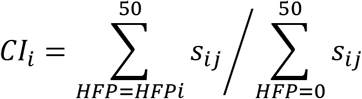

This calculation was repeated for every terrestrial grid cell globally to derive a spatial map of contextual intactness for each taxonomic group (vertebrates, invertebrates, plants). The spatial layers for these three taxonomic groups were then averaged to derive a single contextual intactness layer for biodiversity (Fig. 2).

### Protected area assessment

To determine the coverage of high-value habitat within protected areas, we first identified high-value intact habitat as those grid cells with contextual intactness value greater than 0.5. These are locations which are in a better condition (lower human footprint) than more than half of the similar habitat. We rasterized data from the World Database on Protected Areas (*44*) extracted in August 2018, and converted to a binary raster (protected, not protected) following Butchart *et al*. (*45*). This involved excluding those internationally designated sites not considered as protected areas, excluding ‘proposed’ sites and those with an unknown status, represented sites without a defined shape as geodetic buffers of the appropriate area, and excluding marine-only sites as well as the marine portion of coastal sites. Based on this rasterized protected area layer, we calculated the proportion of high-value habitat within protected areas (Fig. S3).

## Acknowledgments

We thank Sam Nicol and Sam Andrew for providing comments on an earlier version of this work. This research was funded collaboratively by Wildlife Conservation Society – University of Queensland and CSIRO Land and Water. All authors contributed to conceiving the study and designing the analyses. K.M., T.D.H. and C.W. performed the analyses. All authors contributed to writing the manuscript.

## Supporting Information

**Fig. S1.**
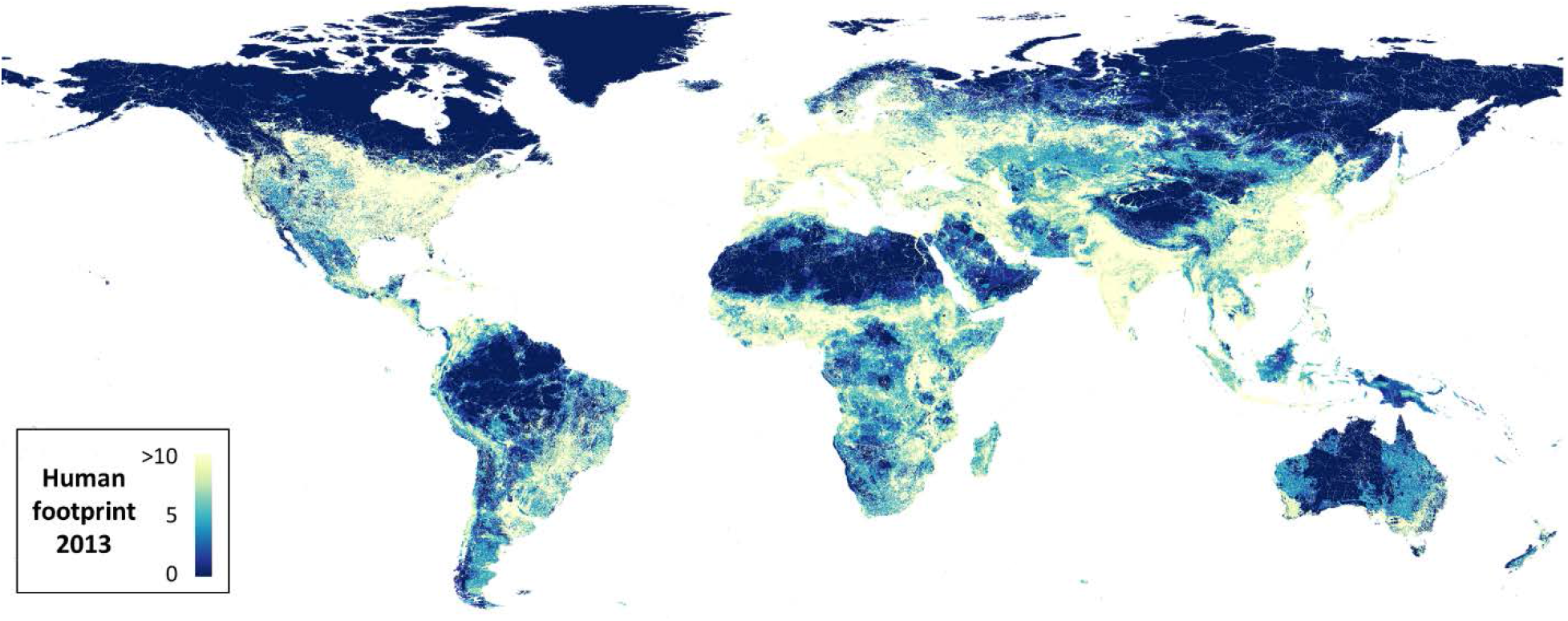
The human footprint on the terrestrial environment, as applied in the present analysis. This layer was developed as described in Venter *et al*. (*21*) but incorporating updated datasets centered on the year 2013. Note that the color ramp scales from 0 to 10, with all human footprint values greater than 10 shown as the same color.

**Fig. S2.**
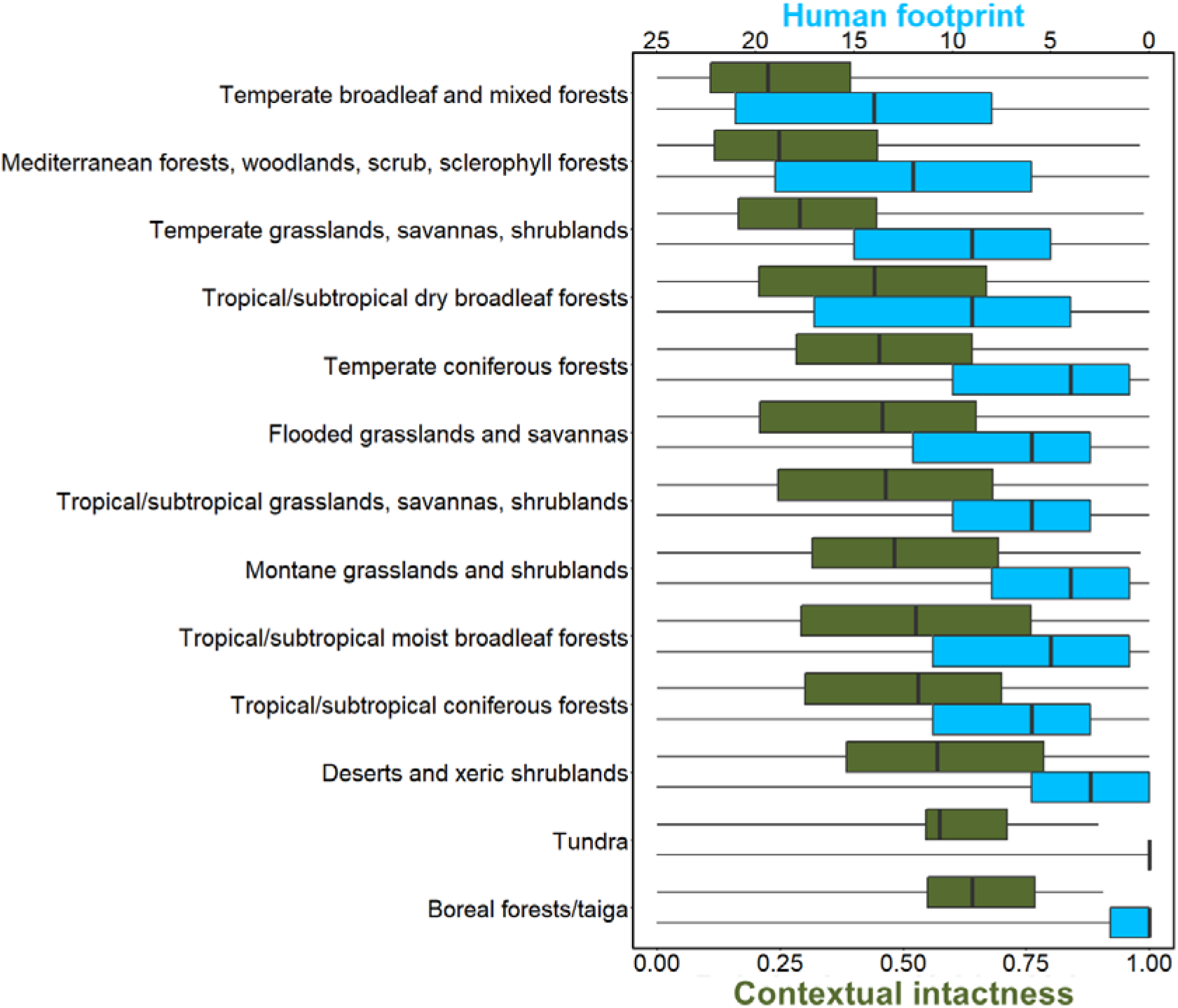
Distribution of contextual intactness and human footprint across the world’s terrestrial biomes. Biomes are ordered based on increasing median contextual intactness.

**Fig. S3.**
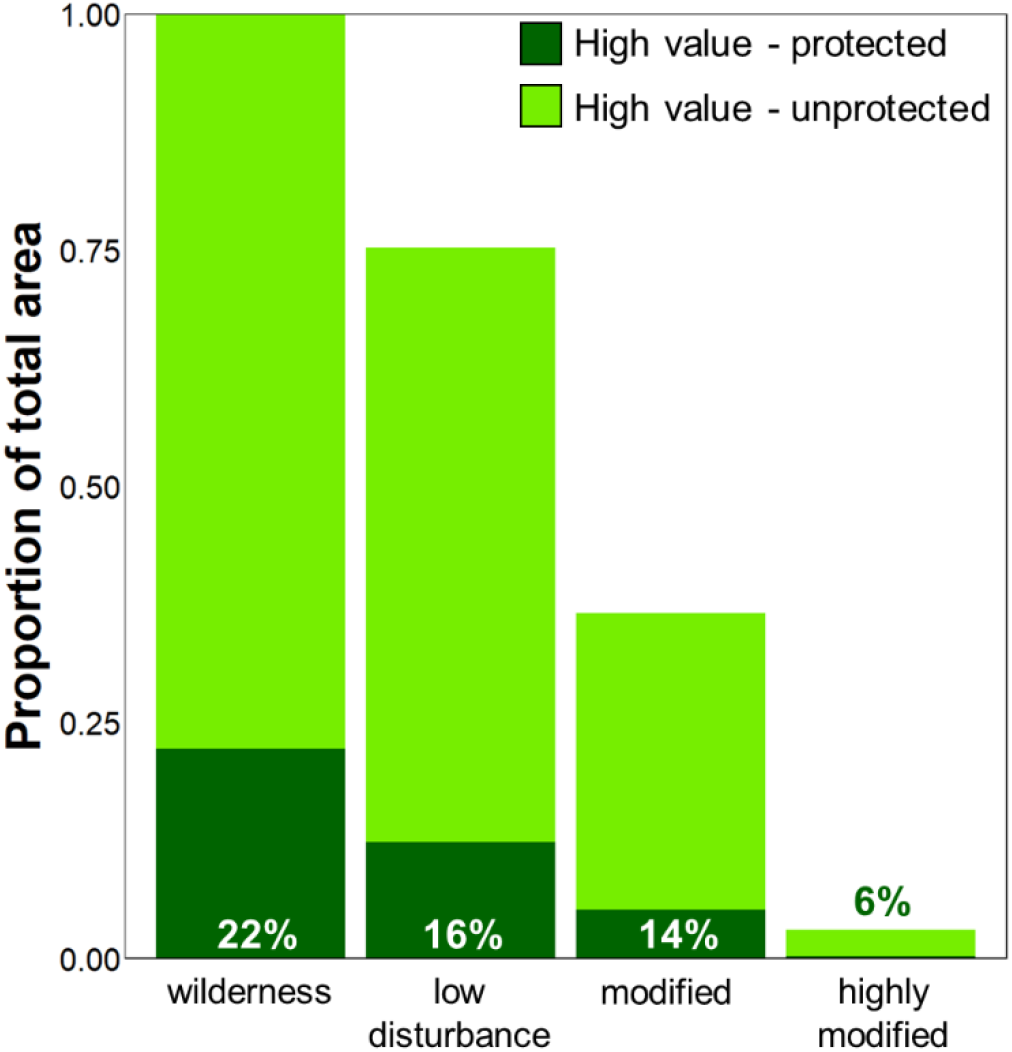
The proportion of high-value habitat (contextual intactness > 0.5) in each of four major levels of human footprint. Also shown is the amount of high-value habitat within protected areas, including the percentage of that coverage.

